# Antiviral Antibody Epitope Selection is a Heritable Trait

**DOI:** 10.1101/2021.03.25.436790

**Authors:** Thiagarajan Venkataraman, Cristian Valencia, Massimo Mangino, William Morgenlander, Steven J. Clipman, Thomas Liechti, Ana Valencia, Paraskevi Christofidou, Tim Spector, Mario Roederer, Priya Duggal, H. Benjamin Larman

## Abstract

There is enormous variability in human immune responses to viral infections. However, the genetic factors that underlie this variability are not well characterized. We used VirScan, a high-throughput viral epitope scanning technology, to analyze the antibody binding specificities of twins and SNP-genotyped individuals. These data were used to estimate the heritability and identify genomic loci associated with antibody epitope selection, response breadth, and the control of Epstein-Barr Virus (EBV) viral load. We identified 4 epitopes of EBV that were heritably targeted, and at least two EBNA-2 binding specificities that were associated with variants in the MHC class-II locus. We identified an EBV serosignature that predicted viral load in white blood cells and was associated with genetic variants in the MHC class-I locus. Our study provides a new framework for identifying genes important for pathogen immunity, with specific implications for the genetic architecture of EBV humoral responses and the control of viral load.

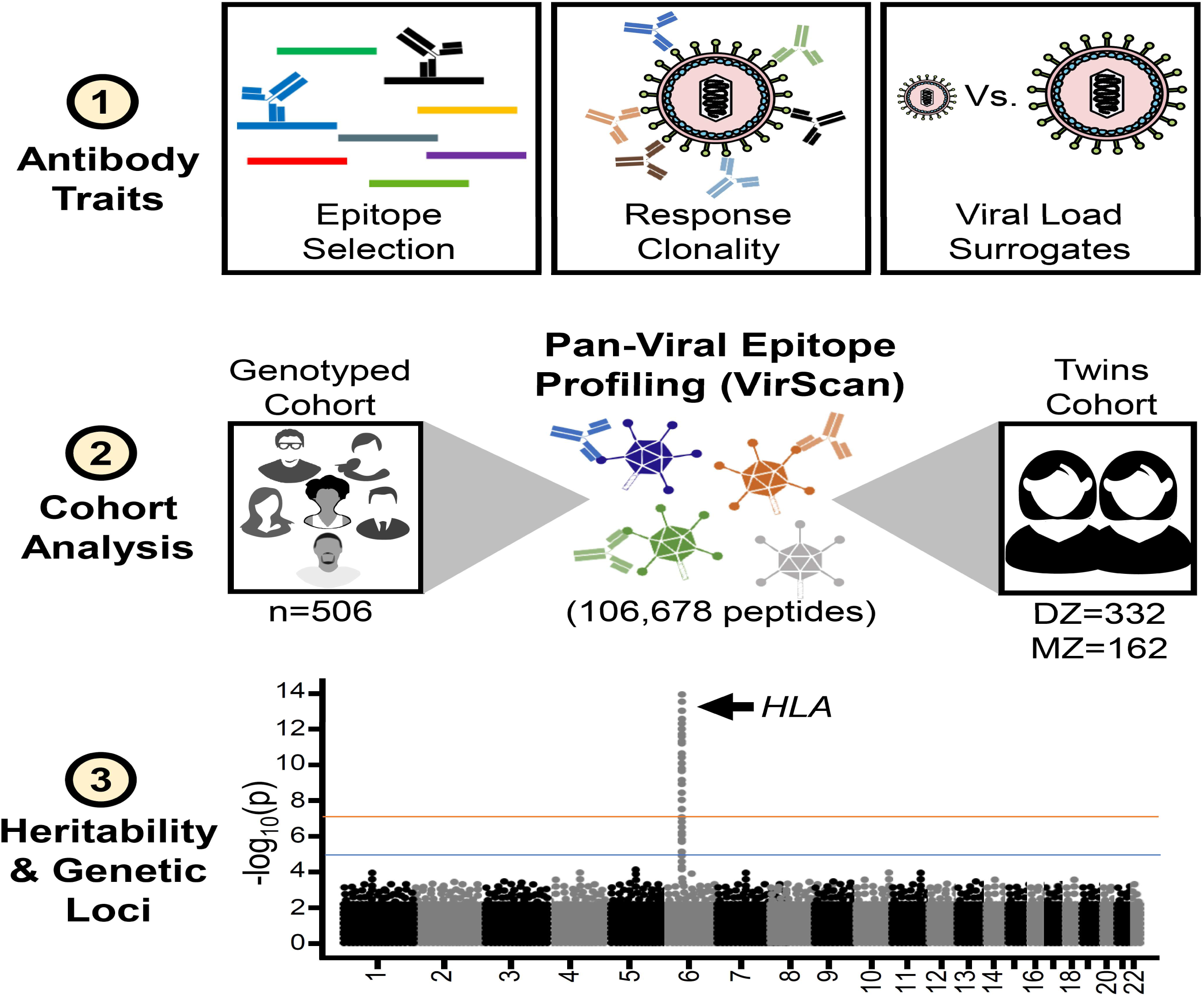

## Introduction

Antiviral antibody responses can last decades after an infection or immunization.^1,2^ They serve as protection from re-infection and can document the exposure history of an individual or population. It has been known for over 50 years that the composition of circulating immunoglobulin (Ig) is influenced by host genetics.^3–6^ Twin, family, and population-based studies have provided examples of heritable contributions to antiviral immune responses; candidate gene and genome-wide association studies (GWAS) have identified genomic loci that influence specific immune traits. However, few studies have examined the heritability of adaptive immune responses broadly across different types of viruses.^7–11^ A recent study investigating the genetic determinants of anti-viral antibody responses (measured by a multiplex serological assay) to 16 common viruses identified strong associations in the HLA locus and in 7 loci outside the HLA.^12^ However, to our knowledge there have been no genetic studies of antibody epitope selection. Antibody fine specificity and breadth (polyclonality) can influence pathogen clearance and protection from re-infection. Genetic variation affecting the expression or function of viral sensing, innate immune signaling, antigen processing and presentation, immune cell function or variation in the antibody locus itself, could all impact the breadth and specificity of an anti-viral antibody response.

Epstein-Barr Virus (EBV) causes a chronic infection with greater than 90% seroprevalence in most adult populations.^13–15^ The EBV genome is relatively large (172 kb), encoding >85 proteins.^16,17^ Primary EBV infections are usually asymptomatic in healthy children and result in a brief episode of infectious mononucleosis in adolescents and older individuals. ^18^ Once the primary infection resolves, EBV establishes a lifelong latent infection residing in circulating memory B-cells.^14^ There is variability in circulating viral load in healthy individuals^19,20^ and the host genes involved in viral load control have not been elucidated. Moreover, Epstein-Barr nuclear antigen 1 (EBNA-1) is an important marker of EBV latency and detection of anti-EBNA-1 antibodies by enzyme-linked immunosorbent assay (ELISA) is often used as an indication of EBV infection. Titers of anti-EBNA-1 antibody in serum of healthy individuals have previously been linked to the HLA-DRB1 and HLA-DQB1 genes from the HLA locus (class II region).^8^ However, the genetic architecture of antibody reactivity to the remainder of the EBV proteome has not be elucidated.

EBV infection can cause nasopharyngeal cancer, Burkitt’s lymphoma, Hodgkin’s lymphoma, and gastric adenocarcinoma, with elevated risk linked to the HLA locus (class I region).^21,22^ Infection with EBV is also believed to play a role in the development of lupus,^23–27^ multiple sclerosis (MS),^28,29^ and other autoimmune diseases, with variants in the HLA locus modulating the risk.^30^ We therefore assessed the genetics of antibody responses against EBV using Phage ImmunoPrecipitation Sequencing (“PhIP-Seq”) with the complete human virome (“VirScan”)^31–33^, in combination with the Anti-Viral Antibody Response Deconvolution Algorithm (AVARDA)^34^. In brief, library scale oligonucleotide synthesis was used to express 106,678 overlapping 56-amino acid peptides spanning all known human viral proteins, in a phage display format, covering ~400 viral species and strains. Phage clones immunoprecipitated by serum antibodies were quantified via Illumina sequencing. We and others have used the VirScan library to characterize the antibody repertoires of preterm neonates^35^, assess viral antibodies after solid organ transplant^36^, to characterize broadly neutralizing HIV antibodies^37^, link enteroviral infection with acute flaccid myelitis^38^, characterize SARS-CoV-2 epitopes^39^, and in large cross-sectional and longitudinal studies of exposure and response to hundreds of human viruses in health and after HIV^40^ or measles infection^41^. Recently, we have advanced the analysis of VirScan data by developing ‘AVARDA’^34^, which calculates likelihoods of viral infections by considering potential cross-reactivities among the peptides and between viruses, and the amount of independent evidence supporting an antibody response to each virus represented in the library. In this study we find that antibody epitope selection is largely a heritable trait, and that class II major histocompatibility (MHC) molecules shape antibody recognition of EBV. We also identified an EBV serosignature that predicts EBV viral load in the periphery and associates with the MHC class I region of HLA.

## Results

### Anti-viral antibody breadth and epitope reactivities are heritable

We used a twins study design to characterize the heritability of VirScan antibody binding specificities. We profiled sera of 494 twins (332 DZ and 162 MZ) from the TwinsUK cohort against the VirScan library. Peptide reactivity scores (z-scores) were calculated by comparing each sample to a set of negative control mock immunoprecipitations (IPs), which did not include serum, and were included on the same plate (**Fig. 1a**).^33,42^

**Figure 1.**
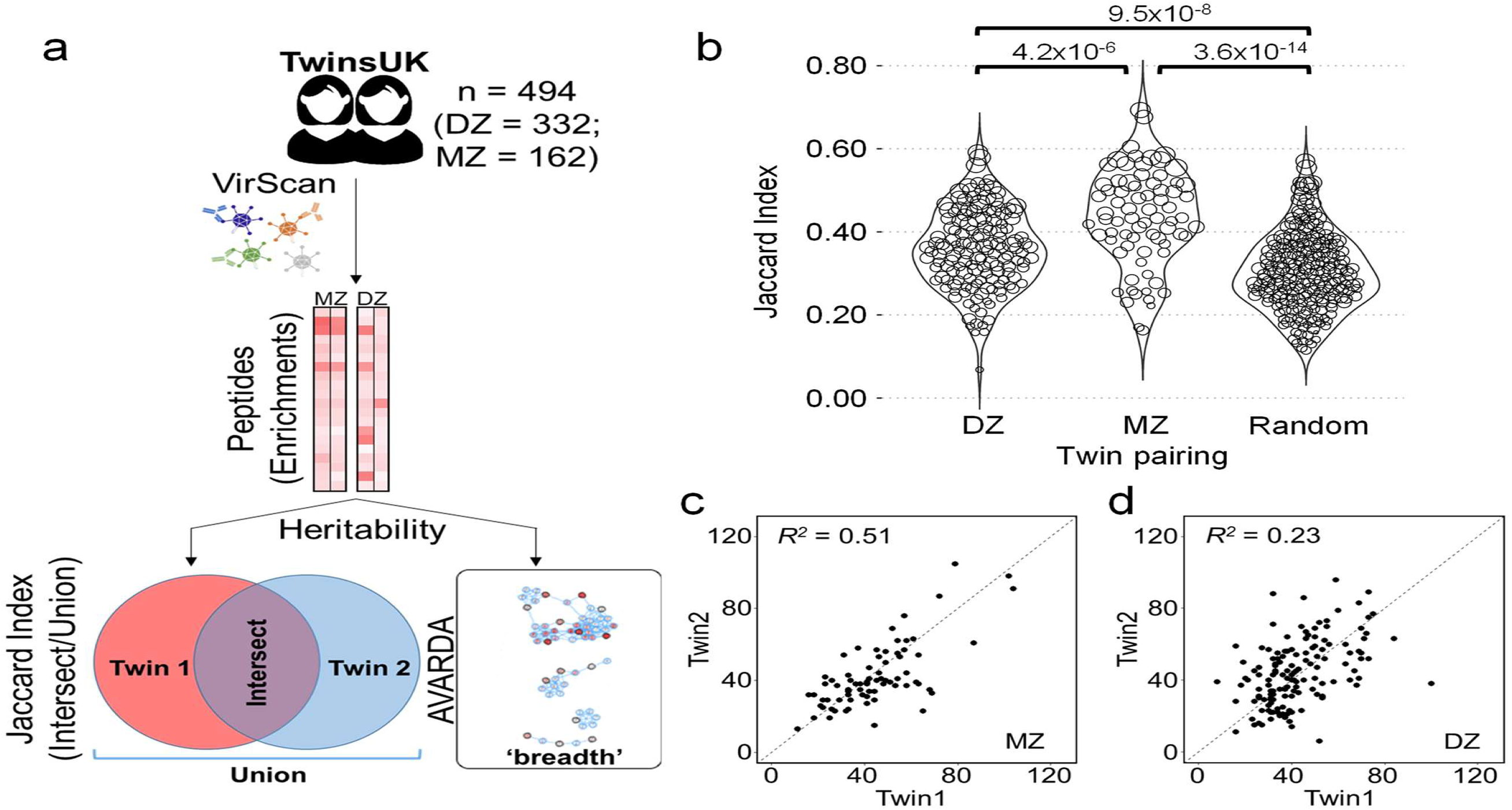
EBV epitope selection and antibody response breadth are heritable traits. **a**, Schematic depicting the heritability analyses. VirScan of the TwinsUK cohort (n = 494; MZ = 81 and DZ = 166 pairs; A upper panel) was used to generate a matrix of peptide reactivity scores. The enrichment scores were used to calculate the Jaccard index and response breadth using AVARDA. **b**, The Jaccard similarity indices of DZ twin pairs (red circles), MZ twin pairs (green circles) or random pairings (blue circles) are shown above. The size of each circle is proportional to the total number of EBV peptides that both twins recognize. **c-d**, Dot plots showing the correlation of EBV response breadth between MZ and DZ twin pairs. Square of the Pearson’s correlation (R^2^) value is provided for each group.

For each virus, we used AVARDA^34^ to calculate the antibody response breadth (polyclonality), and identified seropositive concordant twin pairs. To quantify the contribution of genetic and environmental effects we determined the additive genetic (A), common (C), and unique environmental effects (E) using SEM analysis applied to 446 immunodominant peptide reactivities and the overall breadth of the response to 43 viruses from 9 genera. For each peptide, an individual with a z-score greater than or equal to 7 was classified as a responder and less than or equal to 3 as a non-responder. All values between 3 and 7 were treated as missing data. Only dominant peptides (defined here as reactive in >20% of AVARDA-positive individuals) were included in the analysis; data are presented at the genus level to avoid ambiguity associated with cross-reactivity among viral species with high levels of sequence homology (**Table 1**). The mean breadth values ranged from ~10 peptides up to ~44 peptides with heritability estimates ranging from 5% up to 57%. The virus with the greatest number of dominant peptides, Epstein-Barr Virus (EBV, lymphocryptovirus genus), exhibited heritability ranging from 25% to 89% across 107 peptides.

**Table 1.** Summary of detected viral epitopes and breadth by genus. The table provides a summary of total peptide count, average additive genetic effect (A) component and standard deviation in an ACE model, the mean breadth across all individuals for that genus along with the average A component and standard deviation for breadth. If a genus has multiple species contributing to the data, the mean breadth is marked “X”.

| Virus (Genus) | Peptide Count | Mean Peptide Heritability (A) | Mean Breadth | Mean Breadth Heritability (A) |
| --- | --- | --- | --- | --- |
| Lymphocryptovirus | 107 | 65.3±14.8% | 43.4±17.3 | 39.0% |
| Enterovirus | 56 | 43.9±14.8% | X | 25.8±19.0% |
| Cytomegalovirus | 21 | 62.6±17.0% | 41.3±14.6 | 57.0% |
| Mastadenovirus | 19 | 56.5±14.6% | X | 16.5±13.4% |
| Pneumovirus | 16 | 57.4±20.2% | X | 24.7±1.5% |
| Simplexvirus | 10 | 60.2±13.7% | 36.6±15.6 | 0.0% |
| Alphacoronavirus | 4 | 45.3±27.0% | X | X |
| Betacoronavirus | 3 | 35.1±14.0% | X | X |
| Roseolovirus | 3 | 59.7±14.6% | X | 32.0±15.6% |
| Varicellovirus | 2 | 44.0±19.8% | 10.0±2.9 | 0.0% |

### EBV breadth and epitope selection are heritable traits

We also performed VirScan analysis of serum from a cohort comprised of 506 community volunteers who were also SNP genotyped, of which 388 were of European ancestry (EUR). **Fig. S1 and S2** compare the antibody reactivities of the TwinsUK and the VRC cohorts against the 2,180 EBV peptides in the VirScan library. Immunodominant reactivities were largely restricted to specific regions of the EBV genome (**Fig. S1a-b**) and were mostly localized to the EBNA family of proteins, the transcriptional activator BZLF1 and the envelope glycoprotein BLLF1 (**Fig. S1c**). About 2,000 sub-dominant responses (present in less than 20% of the cohort) were distributed across the EBV genome (**Fig. S2a-b**), with the greatest number of reactivities against the EBNA proteins, BZLF1, BLLF1, the lytic factor LF3 and latent membrane protein 1 (LMP1) (**Fig. S2c**). These results illustrate that while there are shared features of the anti-EBV antibody response between individuals, there is also great inter-individual variability.

In EBV seropositive individuals, we observed an average of 129 reactive peptides (~6% of the 2,180 EBV peptides in the VirScan library), with a range of 12 to 324. Among these reactive peptides, we examined the similarity of the anti-EBV profiles between twin pairs using the Jaccard index (**Fig. 1b**). The similarity of the response among MZ twin pairs was significantly higher compared to their DZ counterpart (p=4.2×10^−6^, much higher than random pairings, p=3.6×10^−14^), illustrating that host genetics sculpts the repertoire of anti-EBV antibodies.

EBV breadth values among twin pairs, estimated by AVARDA, are shown in **Fig. 1c-d**. MZ twin pairs exhibit a higher level of breadth correlation (R^2^ = 0.51) versus DZ twin pairs (R^2^ = 0.23), indicating that in addition to epitope selection, the total breadth of the anti-EBV antibody response is also a heritable trait. Using SEM, we estimated an additive genetic contribution of 39%, shared environmental contribution of 27% and unique environmental contribution of 34%. (**Table 1**). The stochastic elements of antibody responses are captured in the unique environmental component of this model.

We developed a set of selection criteria to identify candidate peptide reactivities that were heritable in the TwinsUK cohort and were adequately powered in the VRC cohort for GWAS. Of all EBV peptides in the VirScan library, we first selected 144 peptides that were dominant in the TwinsUK cohort (**Fig. S3, boxes 1-3**). The most frequently recognized peptides belong to the EBV nuclear antigen (EBNA) family of proteins, with EBNA-3 and EBNA-2 representing of the most immunogenic peptides (**Fig. 2a-b**). Of the 144 immunodominant peptide reactivities analyzed, 107 (at least 1 peptide from each protein) had an estimated heritability of ≥ 20% (**Fig. 2c; Fig. S3, box 4**) and 38 peptide reactivities were also influenced (at least 3%; mean = 42%) by common environmental factors (**Fig. 2d**). Of the 107 peptides with heritable reactivity in the TwinsUK cohort, 57 were dominant in the VRC/EUR cohort and thus selected for GWAS analysis (**Fig. S3, box 5**).

**Figure 2.**
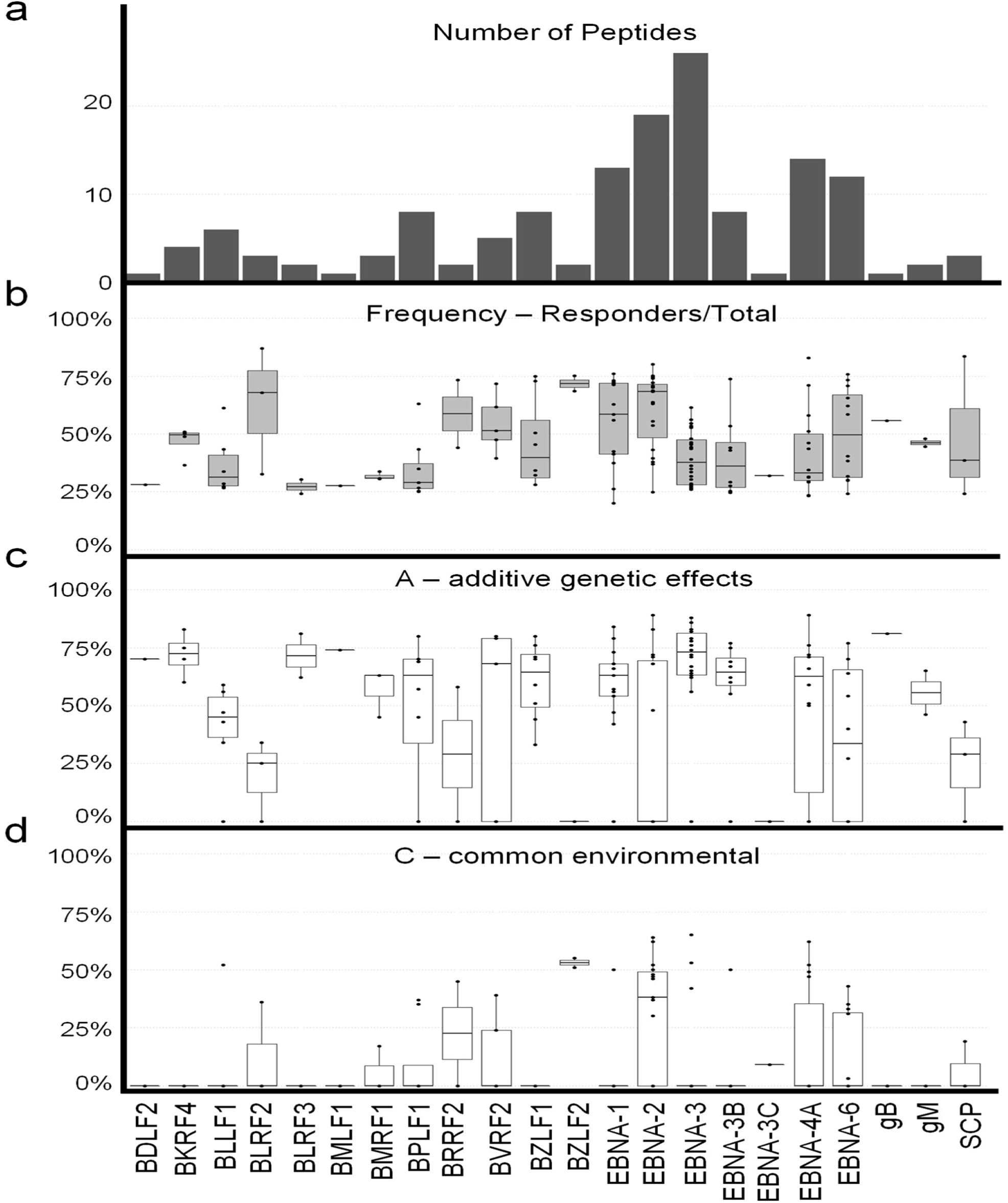
Heritability estimates of individual EBV peptide responses in the TwinsUK cohort. **a**, The number of peptides for each EBV protein associated with dominant antibody responses (at least 20% of the cohort were responders). **b**, Box plot showing the frequency of responders, **c-d**, box plots showing heritable (**c**) and common (**d**) environmental components for peptides from different EBV proteins. Box plots indicate median, interquartile range, and extent of each distribution.

### Genome-wide association studies of EBV peptide reactivities

Single-variant association analyses were performed on the 57 selected peptides and the p-value threshold for significance was adjusted by the Sidak-Nyholt method to account for multiple hypothesis testing (Methods). The Human Leukocyte Antigen (HLA) locus on chromosome 6 was associated (p ≤ 1×10^−9^) with four EBNA2 peptides (**Fig. 3a**). A meta-analysis including the VRC/EUR, VRC/AFR and the TwinsUK cohorts also confirmed the associations in this locus (**Table 2**). These four peptides are in the C-terminal transactivating domain of EBNA-2 (**Fig. 3b**). The magnitude of the antibody response also increased linearly with the number of effect alleles present in the individuals of the VRC/EUR sub-cohort (**Fig. S4a**) and the overall VRC cohort (**Fig. S4b**). The four top associated variants are in linkage disequilibrium (*D’* > 0.8 for each variant pair). There is an overlap in the 95% credible intervals for the four EBNA-2 peptides (ranging from 50 bp to 200.6 kb), and it is likely that this narrowed region harbors the causal variant(s) explaining the differential reactivity of the four peptides (**Fig. 3c-f**). A search of the GWAS catalog showed that the same potentially causal variants identified in this study have also been associated with several different diseases and phenotypes, some with a known or suspected role for EBV infection, including Hodgkin’s lymphoma^43^ and ulcerative colitis^44^. A summary of these associations is provided in **Table S1**.

**Figure 3.**
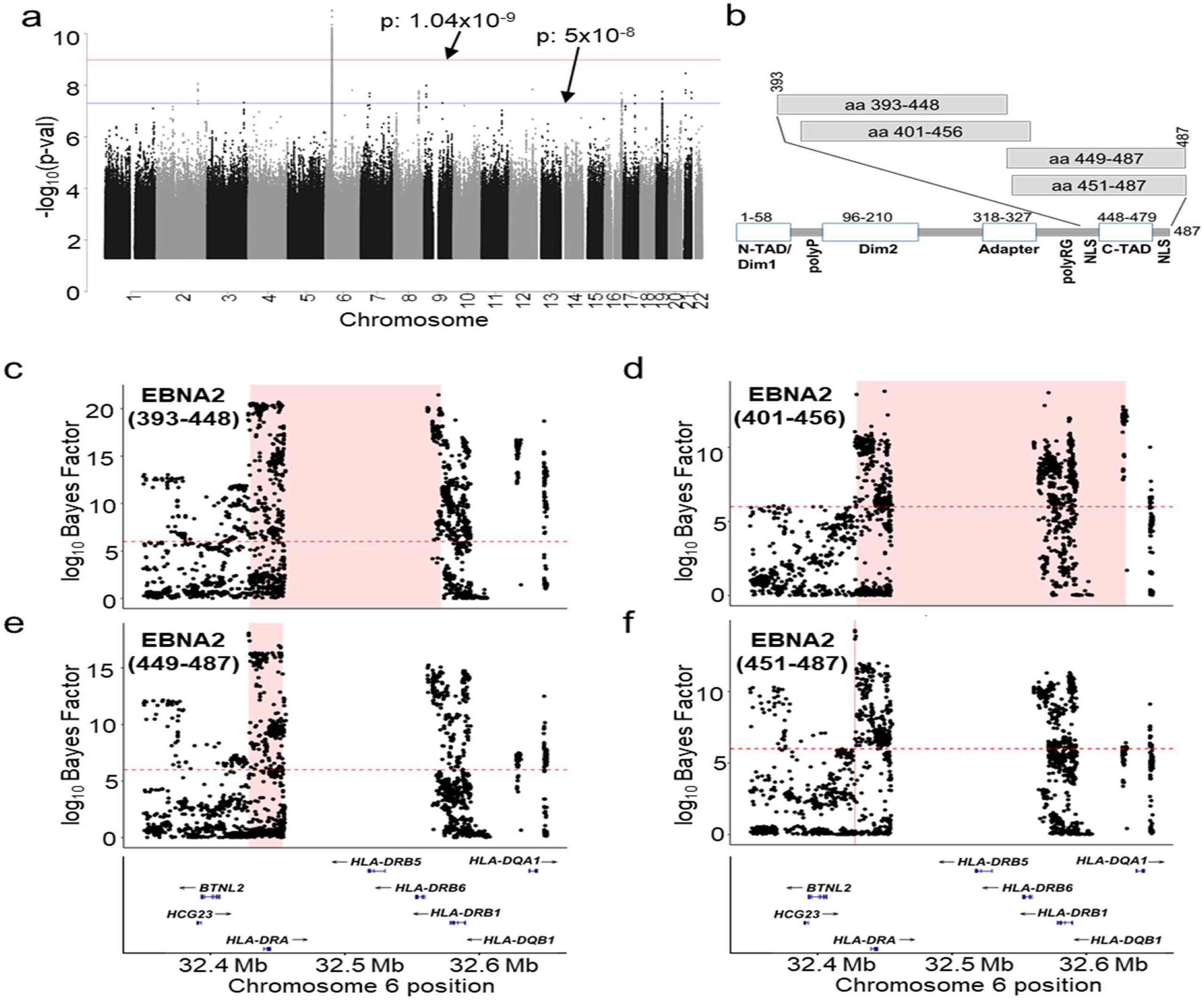
Antibody responses against specific EBV peptides associate with the HLA locus. **a**, The meta-manhattan plot shows a summary of all associations for 57 serodominant peptides in the individuals of European descent in the VRC cohort showing a strong peak on chromosome 6 corresponding to the HLA locus. P-values less than 1.04 × 10^−9^ were considered significant (red line) and those below 5 × 10^−8^ were suggestive (blue line) associations. **b**, A schematic representation of EBNA-2 showing major domains. The four HLA associated peptides fall in the C-TAD spanning aa 393-497. **c-f**, Locus zoom plots for the 4 EBNA-2 peptides. The region in pink was identified by credible sets analysis. All variants with a log10 transformed Bayes factor of >= 6 (dashed red lines) were considered significant.

**Table 2.** Antibody responses against specific EBV peptides are influenced by the MHC class-II locus on chromosome 6. The table lists a set of loci and the EBV peptides associated with them. Locus discovery was performed in individuals of European descent (EUR) in the VRC cohort. A meta-analysis of the three cohorts was also performed with p values provided in the final column.

| Gene | Peptide | SNP | Chr | Pos | ALT | Func. | OR 95%CI | P value | OR.TWINS | P.TWINS | OR.AFR | P.AFR | OR.Meta | P.Meta |
| --- | --- | --- | --- | --- | --- | --- | --- | --- | --- | --- | --- | --- | --- | --- |
| HLA-DRA,HLA-DRB5 | EBNA2 (393-448) | rs9268926 | 6 | 32433067 | G | intergenic | 4.32, 2.73 - 6.84 | 3.75E-10 | 1.29, 1.19 - 1.40 | 4.60E-09 | 1.73, 0.52 - 5.73 | 0.368 | 3.81, 2.46 - 5.91 | 2.05E-09 |
| HLA-DRB1,HLA-DQA1 | EBNA2 (401-456) | rs9271420 | 6 | 32587852 | G | intergenic | 2.89, 2.08 - 4.02 | 2.35E-10 | 1.27, 1.15 - 1.39 | 1.57E-06 | 1.61, 0.86 - 3.02 | 0.132 | 2.89, 2.07 - 4.04 | 3.74E-10 |
| HLA-DRA,HLA-DRB5 | EBNA2 (449-487) | rs9268833 | 6 | 32428062 | T | intergenic | 3.61, 2.49 - 5.23 | 1.28E-11 | 1.19, 1.12 - 1.26 | 2.30E-08 | 2.52, 1.29 - 4.92 | 0.007 | 3.17, 2.23 - 4.53 | 1.58E-10 |
| HLA-DRA,HLA-DRB5 | EBNA2 (451-487) | rs2395196 | 6 | 32450154 | T | intergenic | 3.02, 2.05 - 4.44 | 2.05E-08 | 1.25, 1.14 - 1.38 | 4.98E-06 | 2.63, 1.21 - 5.71 | 0.014 | 2.94, 2.07 - 4.17 | 1.51E-09 |

### Antibody correlates of EBV viral load

Despite significant inter-individual variability, cell-associated EBV genomic copy number is considered to be relatively stable over time, reported at 1-30 genomes per million PBMCs.^19,20^ In an effort to understand antibody and host genetic factors that control the EBV copy number set point, we measured PBMC-associated genome copies by qPCR (**Fig. 4a**). This assay, established by Tsai et al, is sensitive enough to detect a single copy of EBV genome in 10^5^ PBMCs.^45^ Only about one third of the VRC cohort had a measurable EBV viral load, with a majority of positives having 5 copies or less per 10^5^ PBMCs (**Fig. S5a**). We found no correlation between EBV viral load and other covariates such as age, ancestry, and sex (**Fig. S5b-c**). We performed genome-wide association using viral load data both as a continuous trait and a dichotomous variable (detected or undetected) to identify host genetic factors that control viral load, but no significant associations were identified. This lack of genetic association is likely due to a combination of a lack of statistical power to detect small genetic effects and an inability to detect cell-associated EBV genomes in all but the highest titer individuals.

**Figure 4.**
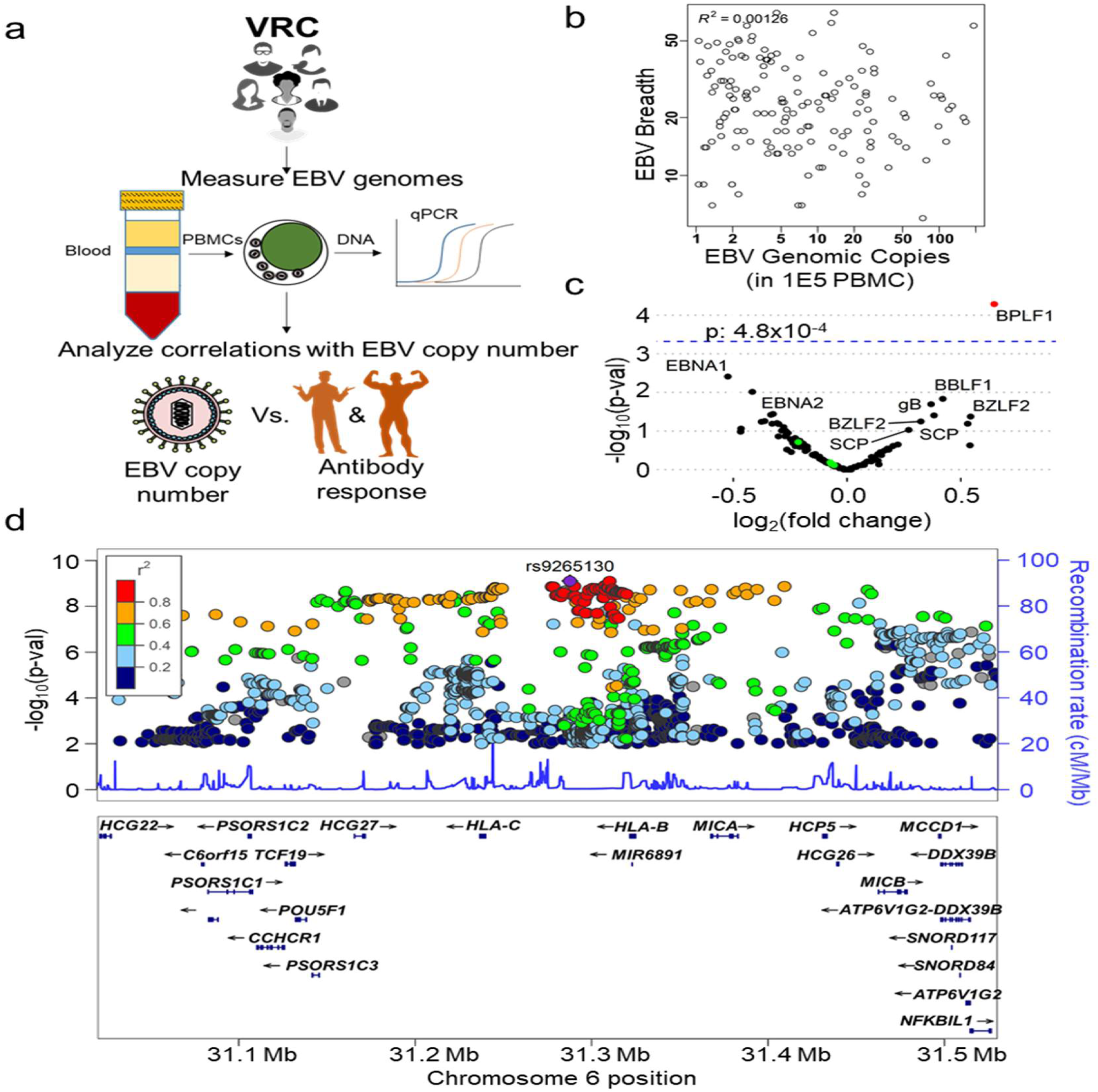
EBV viral load in circulating PBMCs does not correlate with breadth of a response but correlates with responses against specific peptides. **a**, A schematic outlining the approach for EBV copy number measurements. 350ng of DNA (corresponds to roughly 10^5^ cells) extracted from PBMCs was used to detect the presence of EBV genomes by qPCR. Copy numbers were calculated using a standard curve. **b**, EBV copy numbers showed no correlation with breadth of antibody response as calculated by AVARDA. **c**, A volcano plot of significant association between 112 immunodominant EBV peptides (>20% in cohort) and EBV viral load in the VRC cohort. One peptide that showed a significant association (adjusted p-value of 4.8 x 10^−4^) is marked in red. **d**, An LZ plot showing variants in the MHC class-I region on chromosome 6, associated with predicted EBV viral load.

We next examined antibody correlates of viral load. There was no correlation between viral load and overall EBV antibody breadth (**Fig. 4b**). We posited that specific antibody reactivities may be associated with viral load for three reasons: (i) they may directly limit viral replication, (ii) high viral load may stimulate antibody production, and/or (iii) antibodies may correlate with innate immune responses or cell-mediated immunity. We therefore tested individual peptides for their association with EBV viral load (**Fig. 4c**). A single peptide derived from the large tegument protein BPLF1, showed significant association with EBV viral load (Sidak-Nyholt adjusted p-value≤4.8×10^−4^). While not statistically significant, peptides derived from the EBNA family of proteins tended to be negatively associated with viral load and peptides derived from structural proteins from the tegument, capsid and envelope were positively correlated with viral load. The four EBNA-2 C-TAD peptides that associated with variants in the HLA locus (shown as green dots in **Fig. 4c**) were not associated with viral load or sex (green dots in **Fig. S5 d-f**).

In an effort to increase the number of individuals with ‘detectable’ levels of EBV, and to perform analyses in the TwinsUK cohort for which EBV copy number was not available, we used gradient boosting to developed a multi-peptide ‘sero-signature’ predictive of EBV viral load. The 112 peptides reactive in >20% of the VRC cohort were used for model building. ROC analysis was used to identify an appropriate threshold for predicting whether a sample should be considered EBV positive (high copy) or negative (low copy). We performed a GWAS for predicted EBV copy (high versus low) in both the VRC and the TwinsUK cohorts. Meta-analysis revealed a strong association with variants in the MHC class-I locus of the HLA region (**Fig. 4d**, **Table 3**). Our results support a role for CD8+ cytotoxic T cells, possibly in conjunction with antibody epitope selection, in controlling PBMC-associated EBV copy number set point.

**Table 3.**
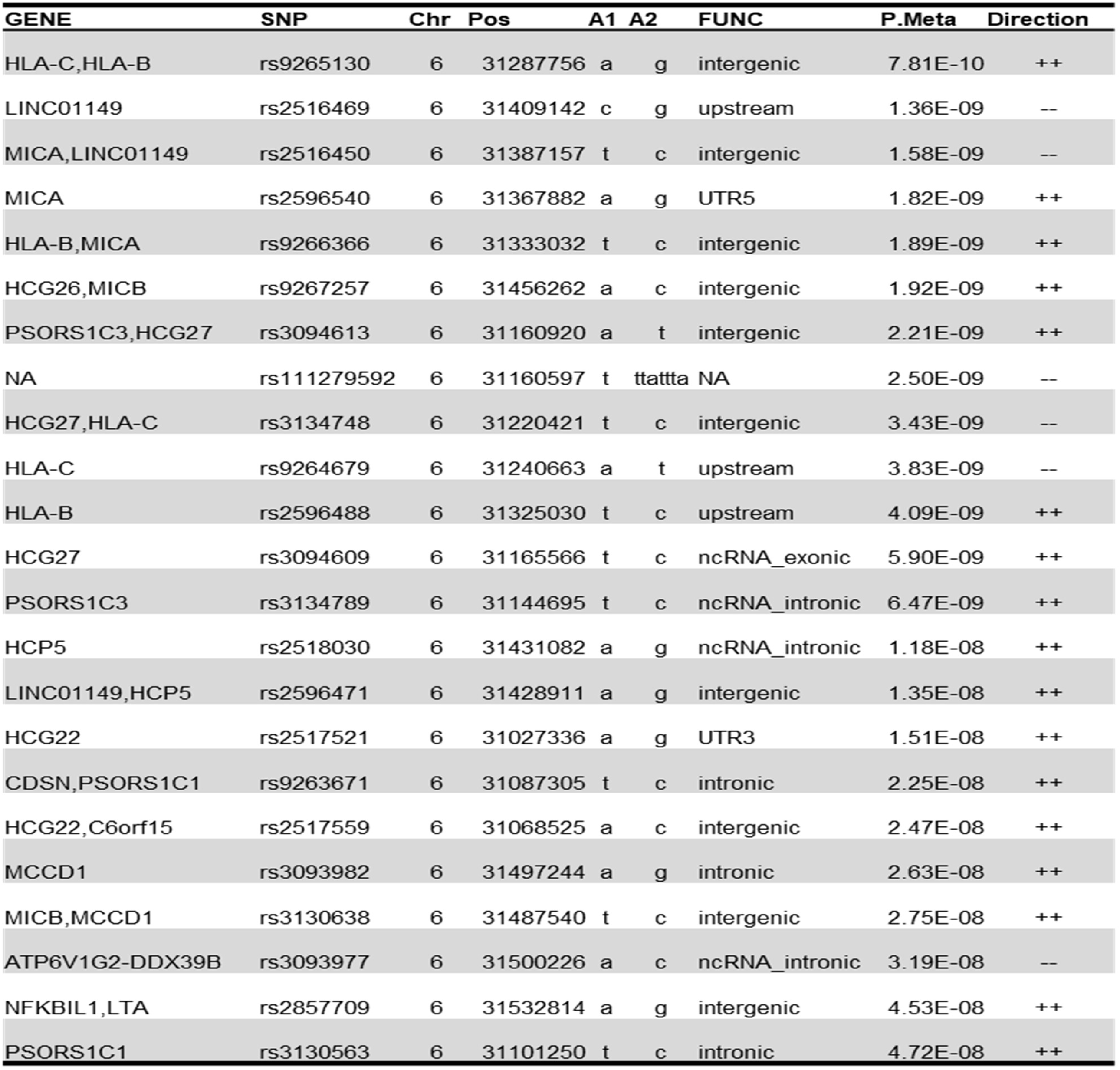
Viral load predictions made from antibody reactivity data are influenced by the MHC class-I locus on chromosome 6. The Table shows a summary of top variants associated with predicted EBV viral load from a GWAS meta analysis on the VRC and TwinsUK cohorts.

## Discussion

The wide variation in human antibody responses to viral infections is influenced by both heritable and non-heritable factors. Previous studies have addressed the role of host genetics by measuring traits such as seroconversion rates and antibody titers to specific antigens.^9,10,46,47^ However, no published studies have yet examined the heritability of antibody responses at the epitope level. In this study, we employed VirScan, a virome-wide antibody profiling technology, to characterize the heritability of epitope selection and identify genetic loci of importance. EBV, a chronic infection with >90% seroprevalence in humans, was used as a model virus in a detailed analysis, which included assessing antibody correlates of viral load. Our results describe a strong heritable component of anti-viral antibody epitope selection, which is associated with specific MHC class II genes, and likely other regions that could not be identified due to study power and the complex genetic architecture of these traits.

GWAS identified MHC class-II associations for four peptides from the C-terminus of EBNA-2. This is similar to previously reported EBNA-1 antibody titer associations with HLA-DRB1 and HLA-DQB1.^48,49^ In our study, reactivity to several immunodominant EBNA-1 peptides were strongly heritable but we did not detect any significant genome wide associations, most likely due to insufficient power.

Variation in the immunoglobulin genes is expected to influence the quality of an antibody response. However, we found no variants in this region associated with EBV epitope selection. The lack of association at the IGH locus may reflect the complexity of genetic variation in this region – including structural rearrangements and copy number variation that are not well captured via SNP based arrays.^50^ However, two reactivities were strongly associated with MHC class-II alleles, which are the major determinants of CD4+helper T cell epitope selection.

Increasing the size of the study population and use of sequencing-based genetic analyses will likely reveal a role for the immunoglobulin loci in epitope selection.

We previously developed a summary statistic to capture the clonality of the antibody response (“breadth”) to a protein or virus. This metric was calculated individually for each virus in the library. The breadth of an antiviral response is likely to correlate with greater protection from re-infection, including potentially heterologous protection from similar viruses.^51^ Using TwinsUK VirScan data, we estimated the genetic heritability of the EBV antibody response breadth to be 39%. However, we detected no significantly associated loci using GWAS. This may be because the breadth of an antibody response is a complex trait, which could not be deconvoluted due to the size of the cohorts in this study.

A GWAS of HIV viral load set point identified roles for the MHC class-I locus and a candidate gene study identified CCR5.^52^ A similar study on EBV viral load has not been performed. We find that PBMC-associated EBV viral load significantly associates with antibody reactivities against a peptide from EBV tegument protein deneddylase, BPLF1, but this should be replicated in an independent cohort. Antibody reactivity against EBV structural proteins tended to be positively correlated with viral load, whereas reactivity against EBV nuclear antigens tended to be negatively correlated with viral load. This could be due to the different spectrum of proteins presented to the immune system in individuals with high versus low viral load, and supported the use of a multi-peptide serosignatures as a surrogate for EBV viral load. We therefore employed machine learning to predict EBV viral load using antibody reactivity of 112 peptides. GWAS linked this EBV serosignature with the MHC class-I locus. Cytotoxic CD8+ T cells play an important role in controlling EBV infection by targeting infected B cells.^53,54^ Specific MHC class I variants could enhance or suppress this activity. An important caveat to this analysis, however, is that hidden variables associated with the serosignature (e.g. a coinfection) may underlie an indirect association with EBV viral load.

There are notable limitations of this study. First, the VirScan library is composed of 56-aa peptides displayed on T7 phage. Conformational, discontinuous, and post-translationally modified epitopes are therefore absent from this study. This limitation is may disproportionately impact surface exposed epitopes. Second, the relatively small size of our cohorts limited the power of the GWAS analyses to detect associations with subdominant epitopes, complex genetic interactions, and antibody responses to less prevalent viruses.

In summary, antibody epitope profiling is a powerful approach for characterizing the genetic architecture of immune responses to viruses and other environmental antigens. This study provides evidence that antibody epitope selection is a heritable trait. Host genetics, in combination with prior immune responses likely explains much of the heterogeneity of an infectious course. GWAS identified a role for MHC class II genes in the selection of antibody epitopes, and MHC class I genes in the maintenance of EBV load. Future studies of larger cohorts will likely identify additional genes important for pathogen immunity.

## Author Contributions

T.V., C.V., M.M., P.D. and H.B.L conceptualized the study; T.V., C.V., M.M., W.M. and A.V. performed the formal analysis; T.V. wrote the original draft; T.V., C.V., M.M., P.D., A.V., W.M., T.L., M.R. and H.B.L reviewed and edited the manuscript; P.D. and H.B.L. supervised the project; H.B.L. acquired funding for the project.

## Acknowledgements

This work was made possible by National Institute of General Medical Sciences (NIGMS) grant R01GM136724 and National Institute of Allergy and Infectious Diseases (NIAID) grant U24AI118633 (H.B.L.). M.R. and T.L. are supported by the intramural research program of the Vaccine Research Center, NIAID, NIH. TwinsUK receives funding from the Wellcome Trust (212904/Z/18/Z) and European Union (H2020 contract #733100). TwinsUK and M.M. are supported by the National Institute for Health Research (NIHR)-funded BioResource, Clinical Research Facility and Biomedical Research Centre based at Guy’s and St Thomas’ NHS Foundation Trust in partnership with King’s College London. P.C. is founded by the European Union (H2020 contract #733100). CV was supported by the Burroughs-Wellcome funded, Maryland: Genetics, Epidemiology and Medicine training program at Johns Hopkins University

We are grateful to Stephen J. Elledge (Harvard Medical School) for generously providing the VirScan library used in this study. We are grateful to the twins who took part in TwinsUK and the whole TwinsUK team, which includes laboratory technicians, administrative staff, and research managers.

## Methods

### PhIP-Seq/VirScan

PhIP-Seq and VirScan have been previously described in detail.^33,55^ Briefly, ELISA was performed to measure total IgG in serum samples and input volume was adjusted to 2 μg of IgG input per IP. VirScan library was mixed with diluted serum at 10^5^-fold coverage (about 9.6×10^9^ pfu for the 56-mer virome library). The library/serum mixture was allowed to rotate overnight at 4 °C, followed by a 4-hour IP with protein A and protein G coated magnetic beads. PCR was performed with primers that flank the displayed peptide inserts. A second round of PCR was performed to add adapters and indexes for Illumina sequencing. Fastq files were aligned to obtain read count values for each peptide in the library, followed by calculation of z-scores as previously described.^42^

### Cohorts

The TwinsUK samples were collected prior to 2017 at King’s College London and the VRC samples at the National Institutes of Health (NIH) Clinical Center under the Vaccine Research Center’s (VRC)/National Institutes of Allergy and Infectious Diseases (NIAID)/NIH protocol “VRC 000: Screening Subjects for HIV Vaccine Research Studies” (NCT00031304) in compliance with NIAID IRB approved procedures. The TwinsUK cohort comprised of 494 individuals: 81 MZ twin pairs and 166 DZ twin pairs; all twins were females of European genetic ancestry an average age of 62 years old. The Vaccine Research Center (VRC) cohort comprised of 535 healthy community volunteers in the greater Baltimore/Washington DC area recruited for multiple studies, of which 388 were of European genetic ancestry (EUR), and 147 of African genetic ancestry (AFR). The VRC cohort included 298 men and 233 women with an average age of 35 years (18-70 years range).

### Jaccard index

Jaccard index calculations were performed by transforming individuals with z-score >= 7 as ‘responders’ and those lesser than that as ‘non-responders’ for each peptide. The total number of EBV peptides where both twins were responders were counted and the Jaccard index for each twin pair was calculated using the formula [*J(A,B)* = (|*A*∩*B*|)/(|*A*∪*B*|)], where A is the set of peptides that twin1 responded to and B is a set of peptides for the corresponding co-twin (twin2).

### Binarization of data

The z-score values from PhIP-Seq were transformed to binarized (response = “1”, non-response = “0” and indeterminate values = “NA”) using a threshold of >=7 as a “1”, <= 3 as a “0” and values between 3 and 7 as “NA”.

### Estimation of infection probability and breadth of a response by AVARDA

A full description of AVARDA is provided in Monaco D. et al.^34^ In brief, AVARDA estimates a conservative assessment of the probability of viral infection using VirScan data. There are three modules in AVARDA, the first of which employs a network graph based on peptide-peptide relationships to define the minimum number of independent antibody specificities (i.e., response breadth) required to completely explain a set of observed peptide reactivities. The breadth values for each individual/virus pair estimated by AVARDA were used in downstream analysis such as the estimation of heritability.

### Real time PCR detection of EBV genomes

Taqman primers for EBV EBNA-1 were synthesized by Integrated DNA technologies (IDT). Sequences for EBNA-1 were reported in Tsai DE et al. and are CGT CTC CCC TTT GGA ATG G (ebna1 forward), GAA ATA ACA GAC AAT GGA CTC CCT TAG (EBV ebna1 reverse) and 6Fam-CCT GGA CCC GGC CCA CAA CC-Tamra (EBV ebna1 probe).^45^ A sensitivity of at least 4 copies per reaction was routinely achieved (**Fig. S6**). Measurement of EBV viral load in the VRC cohort was performed using genomic DNA extracted from PBMCs. 350 ng of DNA (equivalent to ~10^5^ PBMCs) was used per sample in each qPCR reaction. qPCR was performed using PrimeTime Gene Expression Master Mix (Integrated DNA Technologies, Cat. No. 1055772) as per manufacturer’s instructions.

### Testing significance of association between peptide responses and EBV copy number

Fisher’s exact test was used to calculated p-values of association between specific anti-peptide responses and EBV copy numbers measured by real time PCR.

### Peptide selection

We developed a set of criteria to select peptides for GWAS, provided in the flowchart of **Fig. S3**. The VirScan library is composed of 106,678 56-aa peptide sequences representing all known human viruses (~400 viral species and strains, **Fig. S3, box 1**). EBV is represented by 2180 peptides in the VirScan library (**Fig. S3, box 2)**. After binarization of the data, we remove peptides where responder proportions were < 20% and > 80% independently in the TwinsUK cohort (**Fig. S3, box 3**). A total of 144 peptides in TwinsUK were selected for heritability analysis by SEM, resulting in 107 peptides with >= 20% heritability (**Fig. S3, box 4**). Of these peptides, we retained those where the responder proportion in the VRC cohort was between 20% and 80%, resulting in 57 peptides that were used for GWAS (**Fig. S3, box 5**).

### Heritability of epitopes using Structural Equation Models (SEM)

We used the classical twin models to define the influence of genetic and environmental factors on the variance of 134 immunodominant peptide reactivity and the overall breadth of the response to each virus included in this study. Twin studies compare the degree of similarity among monozygotic (MZ) twins, who share 100% of their genetics, and dizygotic (DZ) twins, who like other siblings share on average 50% of their genetics. Under the equal environment assumption (EEA), the variance of the trait/phenotype (P) is explained by latent parameters: P = A +C+E Where “A” represents the additive genetic influence, “C” the common or shared environment between the twin pair and “E” represents the non-shared environment (“E” also includes measurement error).^56^ To estimate the heritability, we used Structural Equation Models (SEM), which utilize observed covariances from both MZ and DZ pairs to establish a causal relationship among the covariances and the latent parameters. We investigated ACE, AE and CE models and used the Akaike’s information criterion (AIC) to select the best fitting model. The model (ACE, AE or CE) with the minimum AIC reflects the best balance between explanatory power and parsimony and was the preferred model. Heritability analyses were performed using the package METs (version 1.2.7.1)^57^ in R (version 4.0.2).

### Genotyping and Imputation

The VRC cohort was genotyped using the Illumina Human Omni 5, GRCh37 (Illumina Inc., San Diego, CA). Quality control steps were performed in Plink 1.9^58,59^ and included removing genetic variants with departure of Hardy-Weinberg equilibrium p-value < 10^−6^, (n=19,227) and missing genotype rate >5% (n=17,891). Participants with missing call rate >5% (n=4), sex inconsistences (n=6) and related individuals based on identity by descent (IBD) estimates (pi_hat >0.2) were also excluded (n=19) from the analysis. Principal component analysis was performed with smartpca from the EIGENSOFT^60^ software package to identify genetic ancestry. In the VRC cohort, 388 individuals clustered with European ancestry population-based samples (88 CEU HapMap samples) and 147 individuals clustered with African ancestry population-based samples (77 YRI HapMap samples). This resulted in 2,783,635 million genetic variants with a minor allele frequency (MAF) ≥ 1%. We then imputed genetic variants using the Michigan imputation server^61^ using the 1000 Genomes Phase 3 reference panel. We removed 15,945,186 variants with low imputation quality (R^2^<0.3) and variants with a MAF<5%. The final genetic dataset contained 7,637,921 variants in 535 individuals. Gene annotation was performed using Annovar^62^ (version date 2017-07-17). Genotyping of the TwinsUK cohort has been described previously in detail.^63^ Briefly, TwinsUK samples were genotyped with a combination of two Illumina arrays (HumanHap300, HumanHap610Q). The normalized intensity data for each array were pooled separately. For each dataset the Illuminus calling algorithm was utilized to assign genotypes. No calls were assigned if an individual’s most likely genotyped was called with less than a posterior probability threshold of 0.95. Validation of pooling was achieved via visual inspection of 100 random SNPs. Finally, intensity cluster plots of significant SNPs were visually inspected for over dispersion biased no calling, and/or erroneous genotype assignment. Before the imputation, the following exclusion criteria were applied to each genotype array. Samples: a) call rate <98%; b) heterozygosity across all SNPs ≥2 s.d. from the sample; c) mean evidence of non-European ancestry; identity errors (assessed by pairwise identity by descent (IBD)). SNPs: a) Hardy-Weinberg p-value<10^−6^ assessed in a set of unrelated samples; b) MAF<1%; c) SNP call rate <97% (SNPs with MAF≥5%) or < 99% (for 1%≤ MAF < 5%).

After the Genotype QC stage, the samples from the two arrays were combined and the imputations were performed using the Michigan Imputation Server^61^ using the 1000 Genomes Phase3 v5 reference panel. After imputation 6,903843 SNPs with MAF > 0.05 and imputation quality R^2^ > 0.3 were included in the analysis.

### Genetic associations of peptide reactivities

We performed single-variant association analysis using dichotomized peptide reactivity data, treating seronegative individuals and borderline reactivities as missing data. Peptides were considered sufficiently powered for analysis if they were reactive in at least 20% but no greater than 80% of the study population and were also heritable in the TwinsUK study (n=107 of 144 peptides; **Fig. S3, box 4**). Based on this criterion, 57 EBV peptides were evaluated in VRC European ancestry individuals (VRC/EUR n=388). Meta-analysis was performed using the TwinsUK cohort (EUR n=494) and VRC study participants of African ancestry (VRC/AFR, n = 147) for 22 peptides with significant (4 peptides) or suggestive (18 peptides) loci (p-value ≤ 5×10^−8^) identified in the VRC European individuals. The significance threshold was estimated with the Sidak-Nyholt method^64,65^, accounting for the number of independent traits (n=48) resulting in a genome-wide significance for EBV of p-value ≤ 1.04×10^−9^. The meta-analysis p-value was set to 0.01 in the replication cohorts, or a joint association p-value less than the VCR European only value.

In the VRC cohort, we interrogated 7,637,921 variants for an association with each of the viral peptides using the penalized quasi-likelihood (PQL) approximation to the GLMM (Breslow and Clayton) implemented in the R package Genesis^66,67^. The African ancestry GWAS included the genetic relationship matrix (GRM PC-Relate) as a fixed effect and 10 principal components as random effects.

The genome-wide association analysis in TwinsUK was performed using a mixed-effects linear model implemented in genome-wide efficient mixed-model association (GEMMA) v 0.98.1. GEMMA is designed for GWAS analysis of family-based data by incorporating pairwise kinship matrix calculated using genotyping data in the mixed-effects linear model to correct relatedness and hidden population stratification.

A total of 22 EBV peptides were included in the meta-analysis using the fixed effect inverse-variance weighted method implemented in METAL.

### Credible sets analysis

We combined ethnic-specific GWAS summary statistics using MANTRA (Meta-Analysis of Trans-ethnic Association Studies)^68^ and used the Bayes factor (BF) to generate credible sets. Briefly, we defined a 1 Mb window (500 kb upstream and 500 kb downstream) from the index variant (lowest Bayes factor), and variants were ranked based on their BF. The posterior probability that this variant is driving the region’s association signal was calculated by dividing the variant’s BF by the sum of the BFs of all variants in the region. The final credible set includes all variants with a cumulative posterior probability sum of 95% in the region.

### GWAS catalog search

We looked at previously reported disease associations of variants of interest using the GWAS catalog version 2020-10-20 (http://www.genome.gov/gwastudies/).

For each GWAS in the discovery group (VRC/EUR) we selected the associated variants included within a credible set and report the disease associations presented in the catalog in **Table S1.**

### Multi-peptide serosignature for EBV viral load prediction

To predict the presence or absence of EBV based on VirScan data, we employed the XGBoost software. (https://xgboost.readthedocs.io/en/latest/) Specifically, we sought to classify samples based on presence or absence of EBV (> 0 copies is present and = 0 copies is absent) based on reactivity to 112 immunodominant EBV peptides. XGBoost leverages ensemble learning and gradient tree boosting to perform regression and/ or classification. Machine learning with XGBoost relies on multiple hyperparameters, including size of terminal nodes in regression/ classification trees, subset of samples used per round, subset of features used per round, number of trees, learning rate, regularization term, and others. To create the prediction model, we first subset the VRC samples into a 90% training set and a 10% validation set. We performed a grid search with 10-fold cross-validation on the 90% validation set to tune model hyperparameters; final hyperparameters were those that maximized the cross-validation AUC and were: max_depth = 2, eta = 0.001, gamma = 2, colsample_bytree = 0.5, min_child_weight = 1, subsample = 0.5. We then applied this gradient boost model to the entire 90% training set, and the performance of the model was determined with the 10% validation set. To prevent overfitting, the iteration with the best cross-validation AUC was used (iteration = 45; **Fig. S7a**). This final model was then used to make predictions of expected likelihood of EBV presence in the TwinsUK cohort. Feature importance (Gain) was calculated using XGBoost.

### Receiver operating characteristics (ROC) analysis of viral load prediction models

ROC analysis was performed using the R package ROCR. The predictions generated by the gradient boost model are a range of values between 0 and 1. The 10 most important features of the prediction model are shown in **Fig. S7b**. We defined an optimal cut off threshold to be the value where the sensitivity and specificity curves intersect (**Fig. S7c-d left panels**). The optimal threshold was determined to be 0.4976 and predicted values above the cutoff are designated EBV positive and below are EBV negative. The model performed at 70% sensitivity and 70% specificity (AUC = 0.776) for the training data and at 60% sensitivity and 60% specificity (AUC = 0.655) for the validation data (**Fig. S7c-d right panels**).

---

**Supplemental Figure 1.**
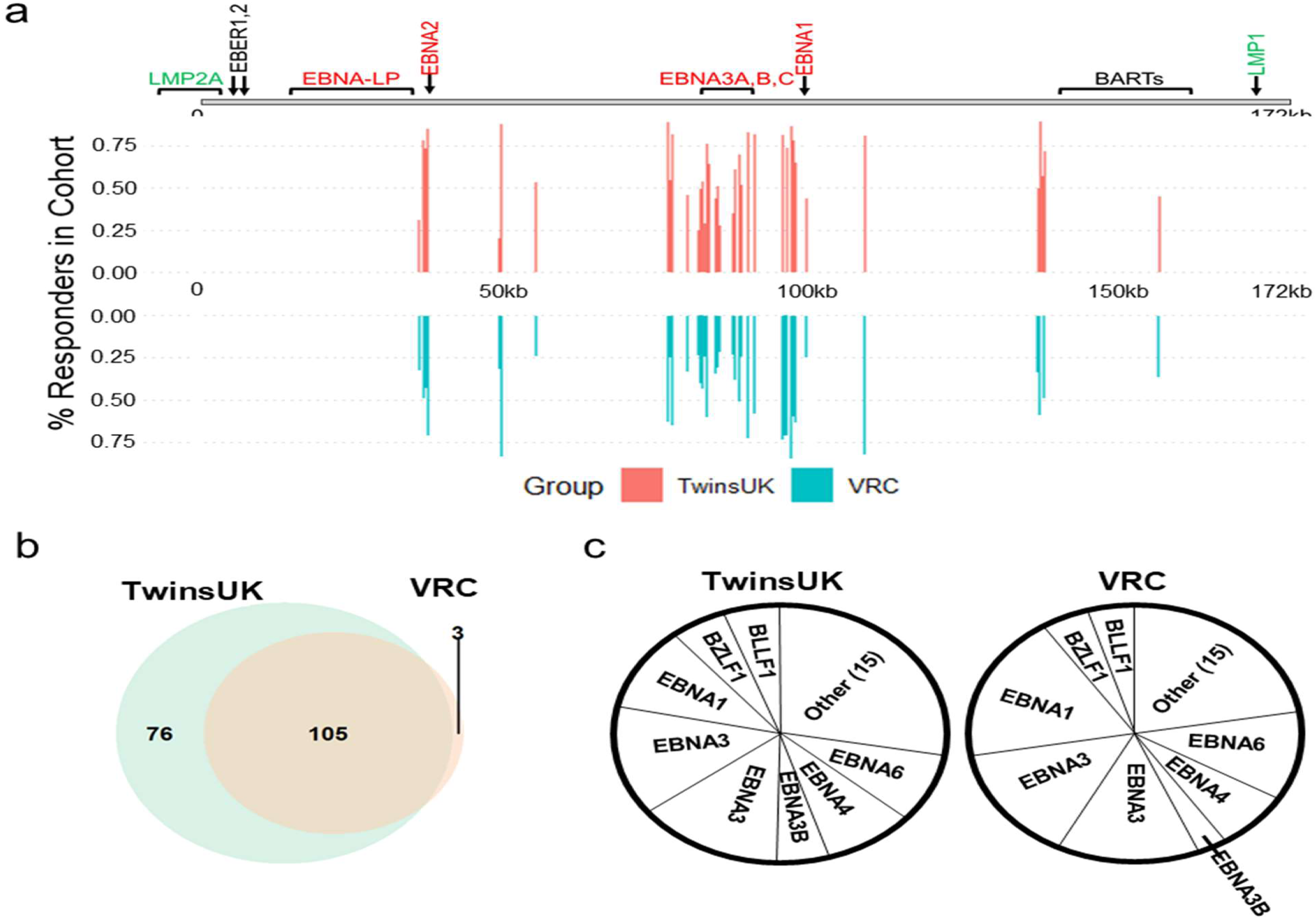
Features of immunodominant anti-EBV antibody responses. **a**, Immunodominant anti-EBV responses (at least 20% of the cohort were responders) shown by genomic position on an EBV reference genome. Genomic map above the plots shows the position of the major nuclear antigens, the LMP proteins and the BART noncoding RNA region. **b**, Venn-diagram of number of immunodominant peptides in both cohorts with 108 peptides in the VRC and 181 peptides in the TwinsUK cohorts. **c**, A piechart of immunodominant responses grouped by protein.

**Supplemental Figure 2.**
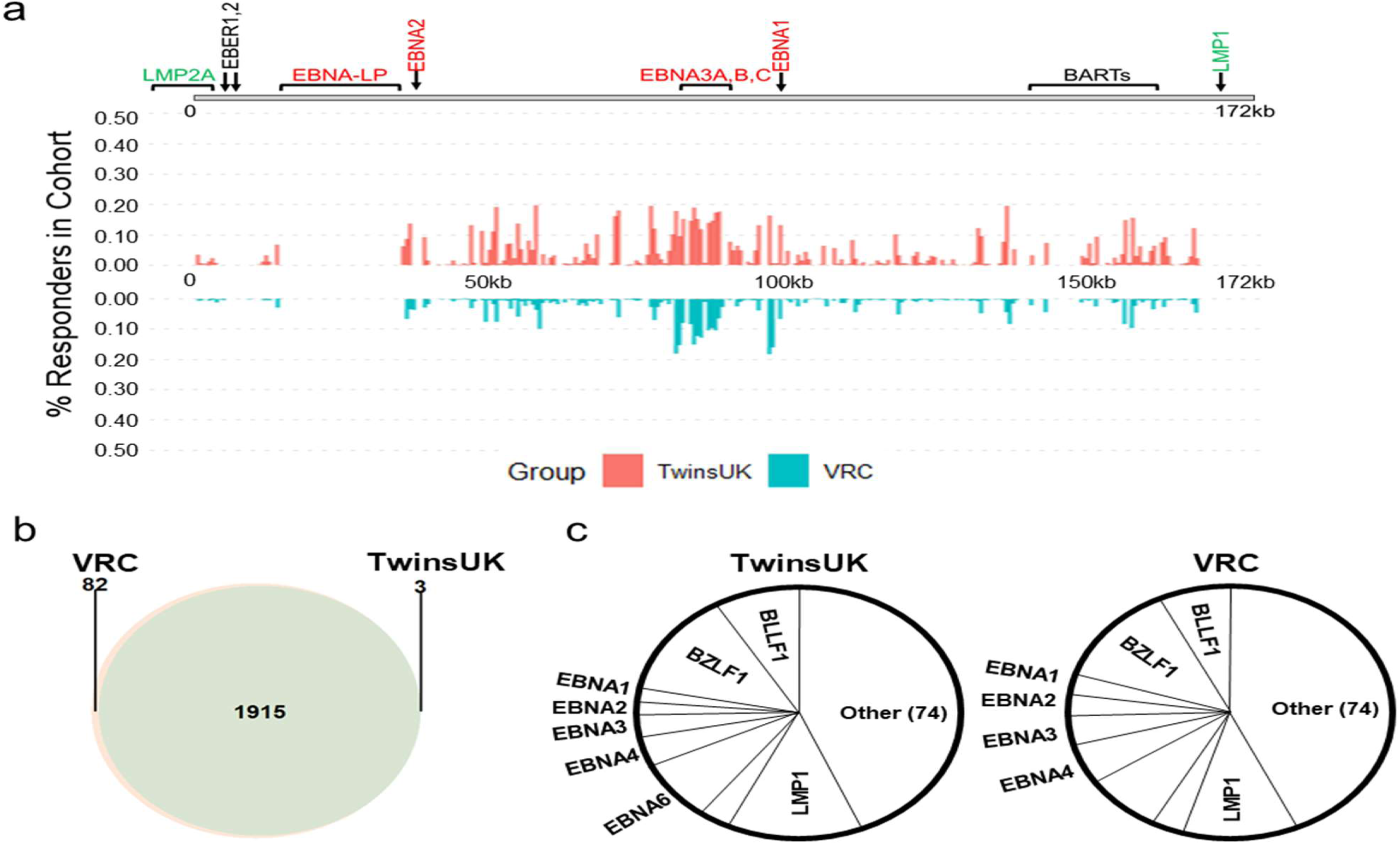
Features of sub-dominant anti-EBV antibody responses. **a**, Sub-dominant anti-EBV responses (less than 20% of the cohort) were seen in peptides that map to most positions on the EBV reference genome **b**, Number of peptides (1997, VRC and 1918 TwinsUK) and **c**, grouping of sub-dominant antibody responses by protein.

**Supplemental Figure 3.**
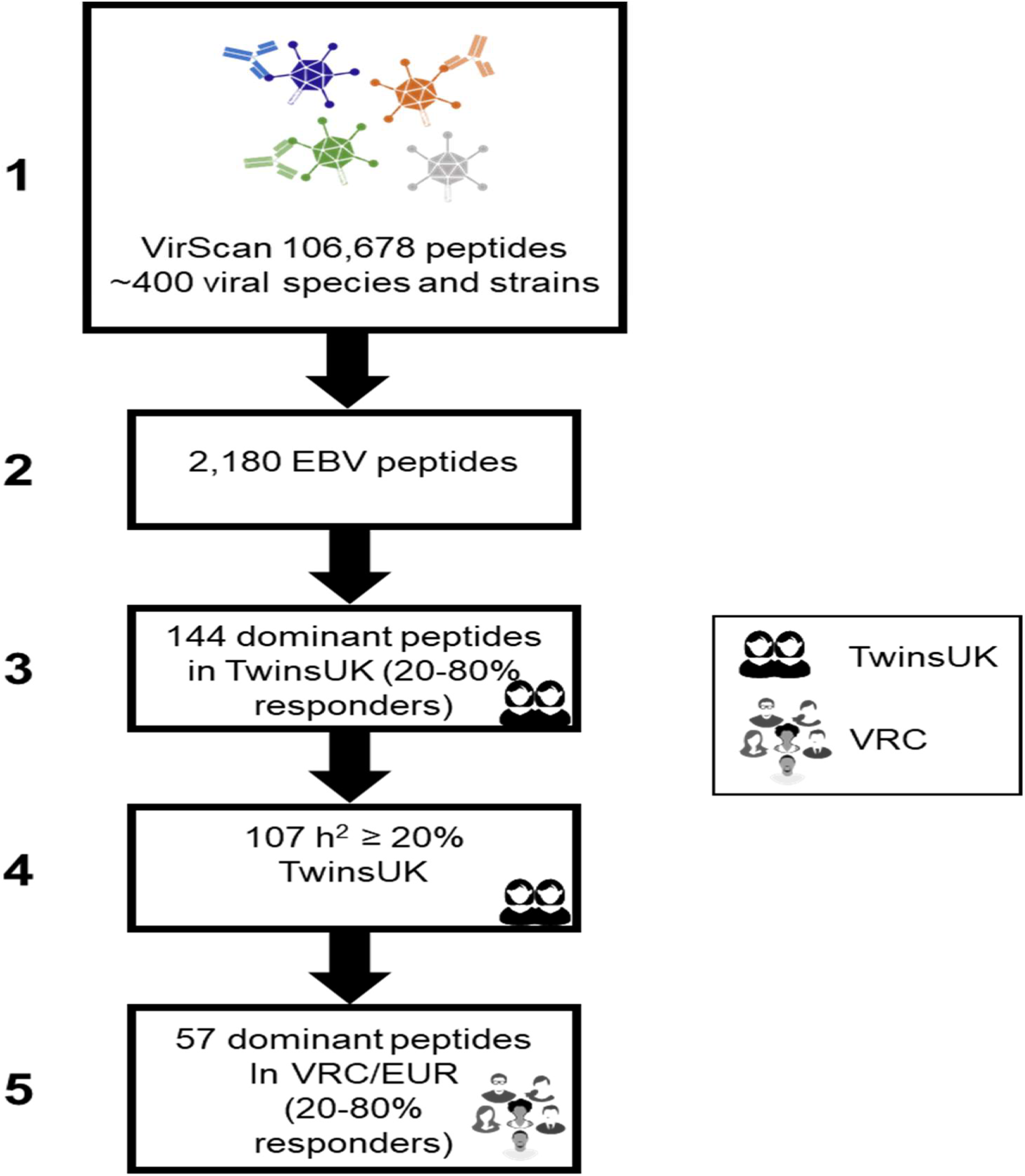
Flow chart of EBV peptide selection for GWAS. Peptide reactivity scores were binarized (Responders: Z-score>=7, non-responders: Z-score <=3) and filtered based on responder proportion (>=20% and <= 80%) in TwinsUK. Of 144 peptides, 107 have estimated heritability >= 20%. A total of 57 EBV peptides were seroprevalent (Responders/non-responders >=20% and <= 80%) in VRC. These 57 peptides were selected for genome-wide association analysis in the VRC/EUR group.

**Supplemental Figure 4.**
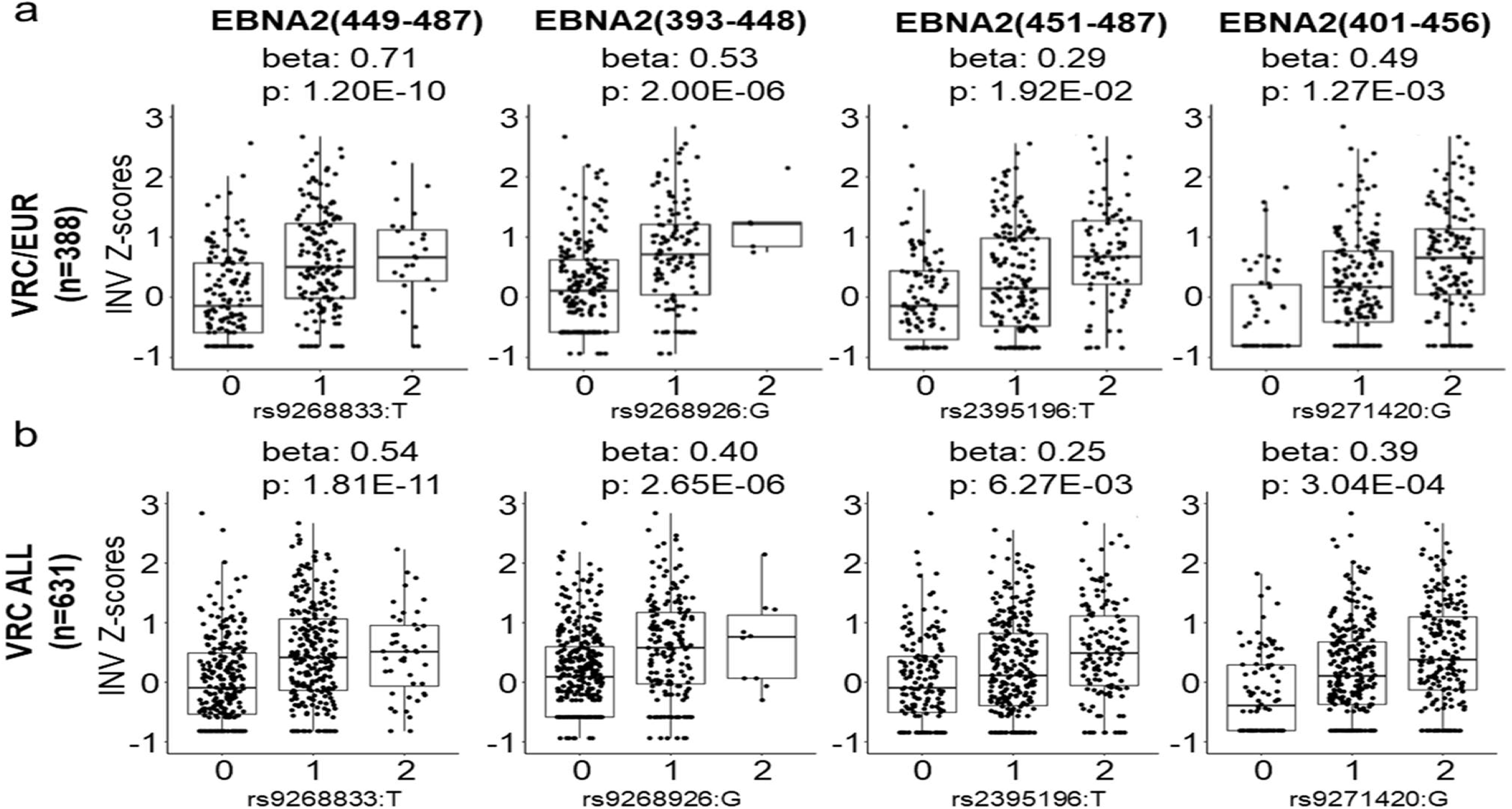
The magnitude of antibody response positively correlates with the number of effect alleles present. **a-b**, Distribution of z-scores in the VRC/EUR sub-cohort (**a**) and the complete VRC cohort (**b**) shows a positive correlation between magnitude of antibody response (higher z-scores equals stronger response) and number of effect alleles in individuals in the VRC cohort.

**Supplemental Table 1.** A list of diseases and phenotypes associated with credible-set variants identified for the two C-terminal EBNA-2 peptides.

| GENE | SNP | Peptide | alt | OR (EUR) | P-Val (EUR) | STRONGEST SNP-RISK ALLELE | DISEASE/TRAIT | OR or BETA | P-Val | PUBMED ID |
| --- | --- | --- | --- | --- | --- | --- | --- | --- | --- | --- |
| HLA-DRB9 | rs2395185 | EBNA-2 (449-487) | T | 1.162 | 3.58E-10 | rs2395185-? | Antinuclear antibody levels | 0.25 | 1.00E-11 | 25186300 |
| HLA-DRB9 | rs28895235 | EBNA-2 (449-487) | A | 1.166 | 3.36E-10 | rs183975233-A | Body mass index | 0.031 | 8.00E-16 | 28892062 |
| HLA-DRB9 | rs9268905 | EBNA-2 (449-487) | C | 1.162 | 3.58E-10 | rs9268905-C | Cystic fibrosis severity | NA | 1.00E-07 | 21602797 |
| HLA-DRB9 | rs9268923 | EBNA-2 (449-487) | T | 1.162 | 3.58E-10 | rs9268923-C | Epstein Barr virus nuclear antigen 1 IgG levels | 0.18 | 1.00E-11 | 26819262 |
| HLA-DRB9 | rs9268923 | EBNA-2 (449-487) | T | 1.162 | 3.58E-10 | rs9268923-C |  | 0.252 | 1.00E-13 | 26819262 |
| HLA-DRB9 | rs9268923 | EBNA-2 (449-487) | T | 1.162 | 3.58E-10 | rs9268923-C |  | 0.225 | 1.00E-11 | 26819262 |
| HLA-DRB9 | rs9268853 | EBNA-2 (449-487) | C | 1.162 | 3.58E-10 | rs9268853-? | Fulminant type 1 diabetes | 3.18 | 2.00E-23 | 30552108 |
| HLA-DRB9 | rs9268905 | EBNA-2 (449-487) | C | 1.162 | 3.58E-10 | rs9268905-? | Giant cell arteritis | 1.79 | 2.00E-54 | 28041642 |
| HLA-DRB9 | rs2395185 | EBNA-2 (449-487) | T | 1.162 | 3.58E-10 | rs2395185-? | Hodgkin's lymphoma | 1.82 | 4.00E-31 | 22286212 |
| HLA-DRB9 | rs2395185 | EBNA-2 (449-487) | T | 1.162 | 3.58E-10 | rs2395185-T | Lung cancer | 1.17 | 1.00E-08 | 23143601 |
| HLA-DRB9 | rs9268853 | EBNA-2 (449-487) | C | 1.162 | 3.58E-10 | rs9268853-C | Lymphoma | 1.56 | 2.00E-10 | 23349640 |
| HLA-DRB9;<br>HLA-DRB5 | rs35957722 | EBNA-2 (449-487) | C | 1.229 | 1.66E-10 | rs35957722-? | Tonsillectomy | 0.0687 | 4.00E-10 | 28928442 |
| HLA-DRB9 | rs9268835 | EBNA-2 (449-487) | A | 1.284 | 1.28E-11 | rs115918645-A | Type 2 diabetes | 1.14 | 1.00E-11 | 29358691 |
| HLA-DRB9 | rs9268835 | EBNA-2 (451-487) | A | 1.168 | 6.13E-10 | rs115918645-A |  | 1.14 | 1.00E-11 | 29358691 |
| HLA-DRB9 | rs2395185 | EBNA-2 (449-487) | T | 1.162 | 3.58E-10 | rs2395185-G | Ulcerative colitis | 1.49 | 9.00E-23 | 20228799 |
| HLA-DRB9 | rs2395185 | EBNA-2 (449-487) | T | 1.162 | 3.58E-10 | rs2395185-G |  | 1.92 | 5.00E-22 | 19915573 |
| HLA-DRB9 | rs2395185 | EBNA-2 (449-487) | T | 1.162 | 3.58E-10 | rs2395185-? |  | 1.52 | 1.00E-16 | 19122664 |
| HLA-DRB9 | rs9268853 | EBNA-2 (449-487) | C | 1.162 | 3.58E-10 | rs9268853-T |  | 1.37 | 3.00E-06 | 23511034 |
| HLA-DRB9 | rs9268853 | EBNA-2 (449-487) | C | 1.162 | 3.58E-10 | rs9268853-T |  | 1.4 | 1.00E-55 | 21297633 |
| HLA-DRB9 | rs9268923 | EBNA-2 (449-487) | T | 1.162 | 3.58E-10 | rs9268923-C |  | 1.45 | 4.00E-15 | 20228798 |
| HLA-DRB9 | rs9268838 | EBNA-2 (449-487) | A | 1.284 | 1.28E-11 | rs114800139-A | Vogt-Koyanagi-Harada syndrome | 3.02 | 1.00E-119 | 25108386 |

**Supplemental Figure 5.**
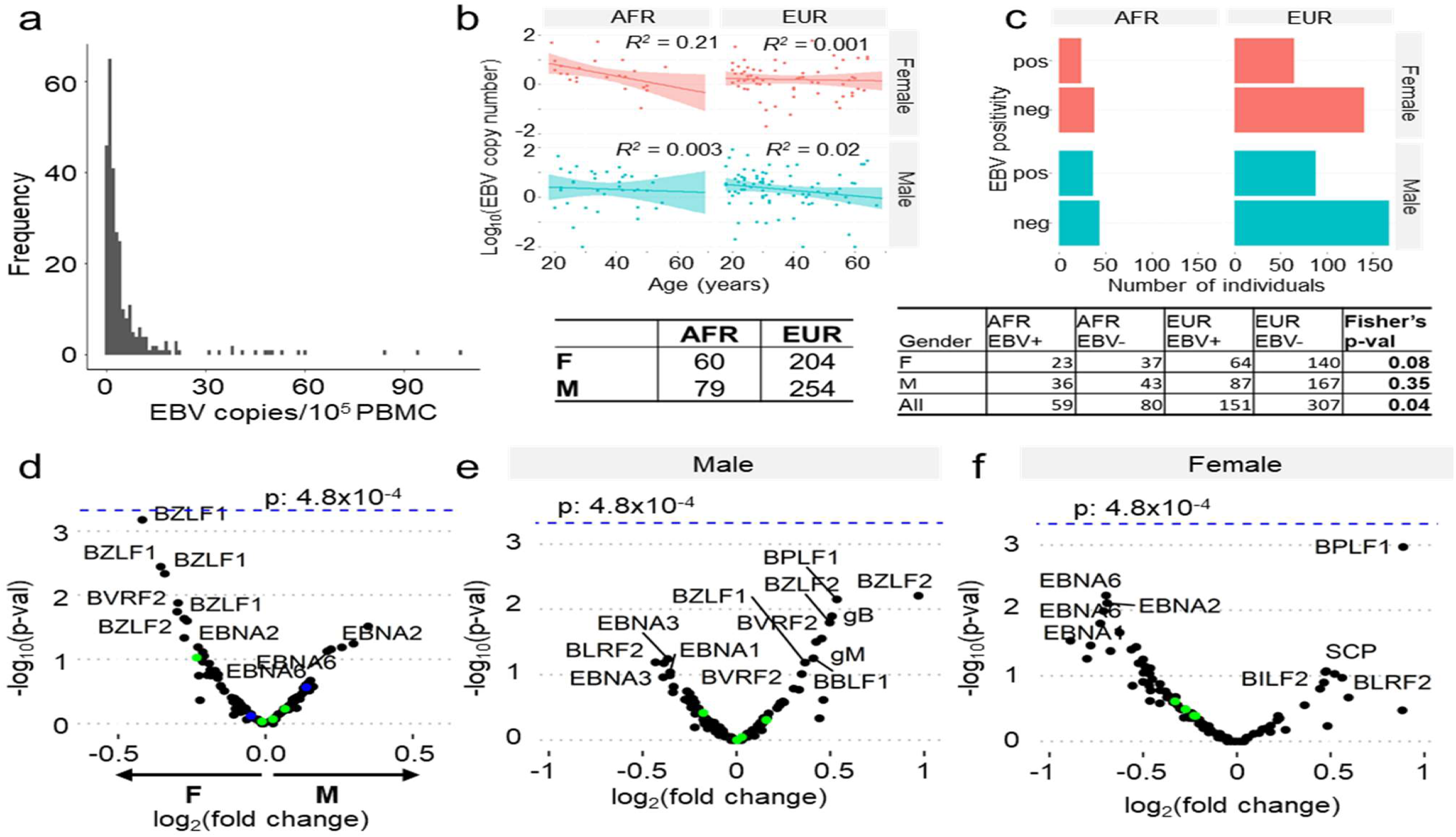
EBV viral load is not correlated with ancestry, age or gender. **a**, A histogram showing the distribution of the EBV genomic copies detected in 10^5^ PBMCs of individuals in the VRC cohort. **b**, Scatter plot of EBV viral load with age, grouped by EUR or AFR ancestry and by gender. **c**, Histogram of EBV positive and negative individuals in the EUR and AFR sub-groups grouped by gender. Tables below the panels provide the number of individuals in each group shown. **d**, A comparison of each peptide specific reactivity between male and female sub-groups in the VRC cohort shows no significant differences. **e-f**, Associations between peptide reactivities and EBV viral load show no differences for men (**e**) or women (**f**).

**Supplemental Figure 6.**
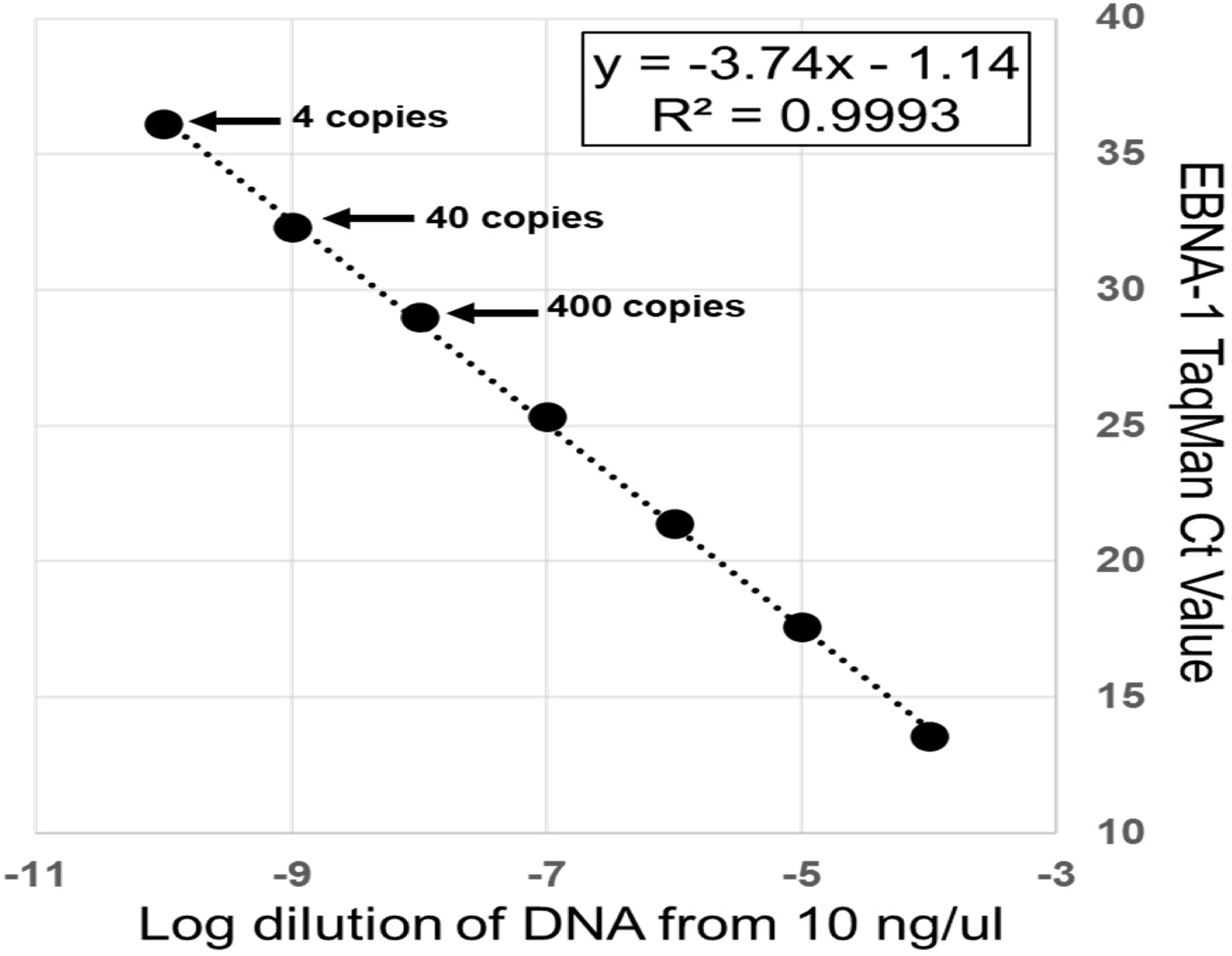
Quantitative PCR detection of EBV EBNA-1 was highly sensitive. The graph shows qPCR performed on a dilution series of EBNA-1 DNA fragment. The assay had a linear range spanning 7 decades and was sensitive to 4 copies per reaction.

**Supplemental Figure 7.**
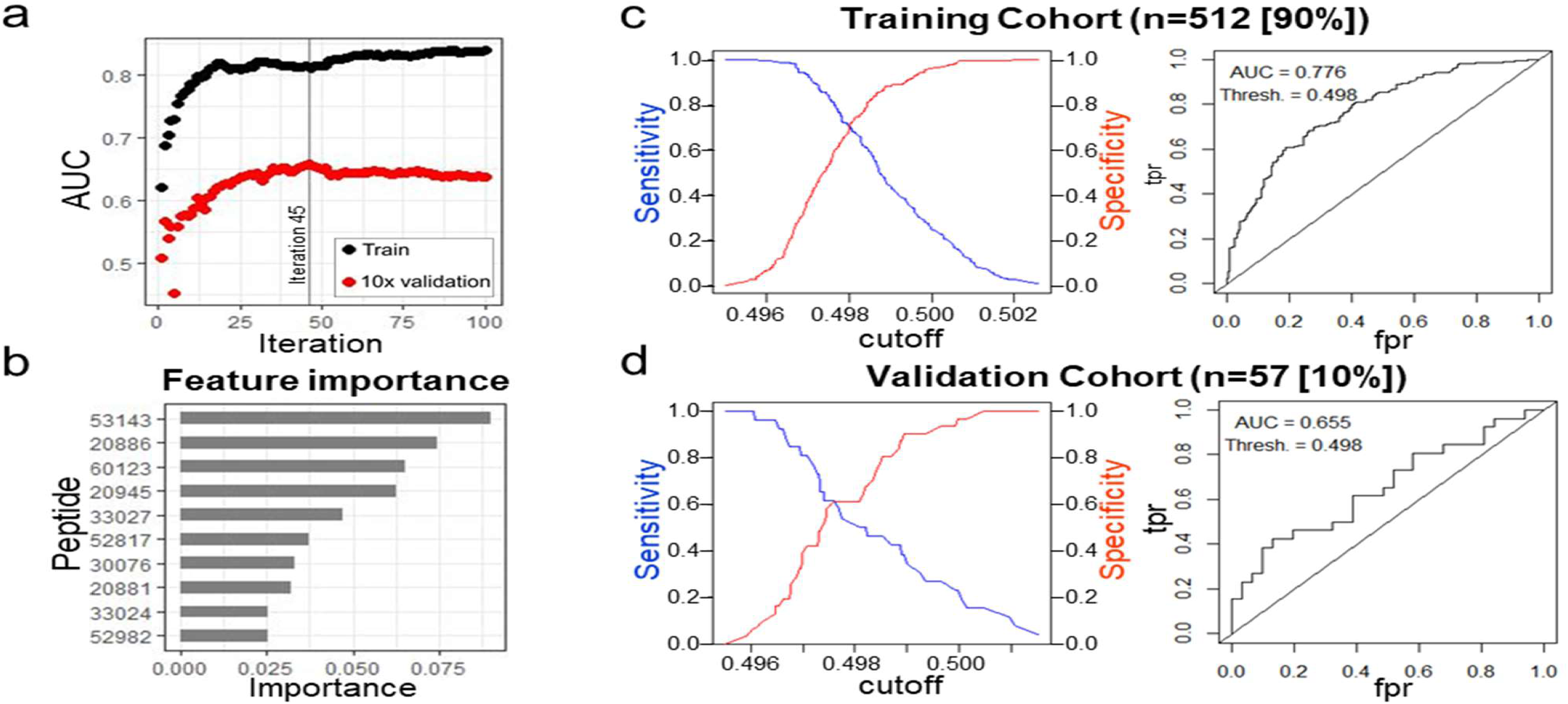
Generation of a prediction model by gradient boosting. **a**, Training of the model. **b**, ROC analysis of training cohort (n = 512, 90%). **c**, The 10 most important features of the prediction model. **d**, ROC analysis of the independent validation cohort (n=57, 10%).

